# PartsGenie: an integrated tool for optimising and sharing synthetic biology parts

**DOI:** 10.1101/189357

**Authors:** Neil Swainston, Mark Dunstan, Adrian J. Jervis, Christopher J. Robinson, Pablo Carbonell, Alan R. Williams, Jean-Loup Faulon, Nigel S. Scrutton, Douglas B. Kell

## Abstract

Synthetic biology is typified by developing novel genetic constructs from the assembly of reusable synthetic DNA parts, which contain one or more features such as promoters, ribosome binding sites, coding sequences and terminators. While repositories of such parts exist to promote their reuse, there is still a need to design novel parts from scratch.

PartsGenie, freely available at http://parts.synbiochem.co.uk, is introduced to facilitate the computational design of such synthetic biology parts. PartsGenie has been designed to bridge the gap between optimisation tools for the design of novel parts, the representation of such parts in community-developed data standards such as Synthetic Biology Open Language (SBOL), and their sharing in journal-recommended data repositories.

Consisting of a drag-and-drop web interface, a number of DNA optimisation algorithms, and an interface to the well-used data repository JBEI ICE, PartsGenie facilitates the design, optimisation and dissemination of reusable synthetic biology parts through a single, integrated application. PartsGenie can therefore be used as a single, stand-alone tool, or integrated into larger synthetic biology pipelines that are being developed in the SYNBIOCHEM centre and elsewhere.

## Introduction

The computational design of synthetic biology parts is greatly aided by the increasing availability of open access tools, standards and data repositories. A number of excellent tools are available to synthetic biologists for designing novel parts, including the RBS Calculator [1] and RedLibs [2] for ribosome binding site (RBS) optimisation, and numerous academic and commercial packages for codon optimisation of coding sequences (CDS) for expression in a given host [3]. In addition to this, the introduction of community-developed standards for representation of synthetic biology designs, such as Synthetic Biology Open Language (SBOL) [4] and SBOL Visual [5] have greatly facilitated the sharing and reuse of synthetic DNA. Indeed, this journal now recommends the sharing of genetic designs, and has a dedicated repository to which synthetic biologists are encouraged to submit synthetic DNA designs to support submitted manuscripts [6].

Despite these advantages, practical problems exist for the synthetic biologist in utilising these packages. First, there is still as yet little integration of tools, such that separate tools are required for RBS optimisation, codon selection, and the optimisation of designs to enable simple (and therefore cost-effective) synthesis by manufacturers. Secondly, tools that support the assembly of synthetic biology parts adhering to data standards [7] typically prioritise the re-use of existing “features”, such as RBSs, CDSs, promoters, terminators, etc., and do not interface with tools that design novel features and designs. Finally, upon submitting designs to manufacturers for synthesis, a further check or optimisation step is typically required to comply with vendor-specific synthesis constraints. Following such a step-wise approach, which is heavily reliant on the use of numerous software packages and manual cut-and-pasting of DNA sequences between tools, is unnecessarily time consuming and potentially prone to user error.

PartsGenie is therefore introduced for the design of synthetic biology parts, integrating many of the above challenges in a single web application, which can subsequently be extended to cover a more comprehensive pipeline for applications such as metabolic engineering. Driven by an intuitive drag-and-drop interface, the user can assemble multiple DNA parts from a range of synthetic DNA features, which can then be sequence optimised simultaneously, with the final results directly exportable to the well-used data repository JBEI ICE [8], following journal recommendations. Such a system, bridging the gaps between multiple design algorithms and standardised data repositories, benefits both the designer in terms of simplification, and the synthetic community as a whole through data sharing.

## Results and Discussion

PartsGenie allows for the flexible design of multiple synthetic biology parts through the use of a simple web interface (Figure 1). Designs can be assembled through the arrangement of multiple features, which can be drag-and-dropped from a palette that include RBSs, CDSs, promoters, etc., represented as glyphs in SBOL visual format [5]. Furthermore, user-defined and randomly generated sequences may also be included, which can represent flanking regions used for assembly, and spacer regions respectively. Designs and individual features can be named, and parameters required for their optimisation can be specified by selecting the feature and filling in the resulting form. Examples of such parameters are a translation initiation rate (TIR) for RBSs, and desired amino acid sequence for CDSs. In specifying amino acid sequences, PartsGenie offers an integrated UniProt search tool, allowing sequences, ids, NCBI Taxonomy terms and enzyme classification (EC) numbers to be automatically extracted and associated with the CDS, facilitating future traceability. Extracted sequences can subsequently be edited directly in the Feature panel, to introduce mutations and truncations etc., if necessary. A host organism is needed for designs that include RBS or CDS features, in order to determine the optimum Shine-Dalgarno sequence for RBSs, and prefered codon usage for CDSs. (Supported prokayotic organisms are selected from a simple autofill field, and the backend algorithm extracts Shine-Delgarno sequences and codon usage tables automatically from the RBS Calculator and Codon Usage Database [9] respectively.) Multiple designs can be submitted as a single optimisation task, and designs may be copied in the interface to facilitate the submission of multiple, related designs.

**Figure 1:**
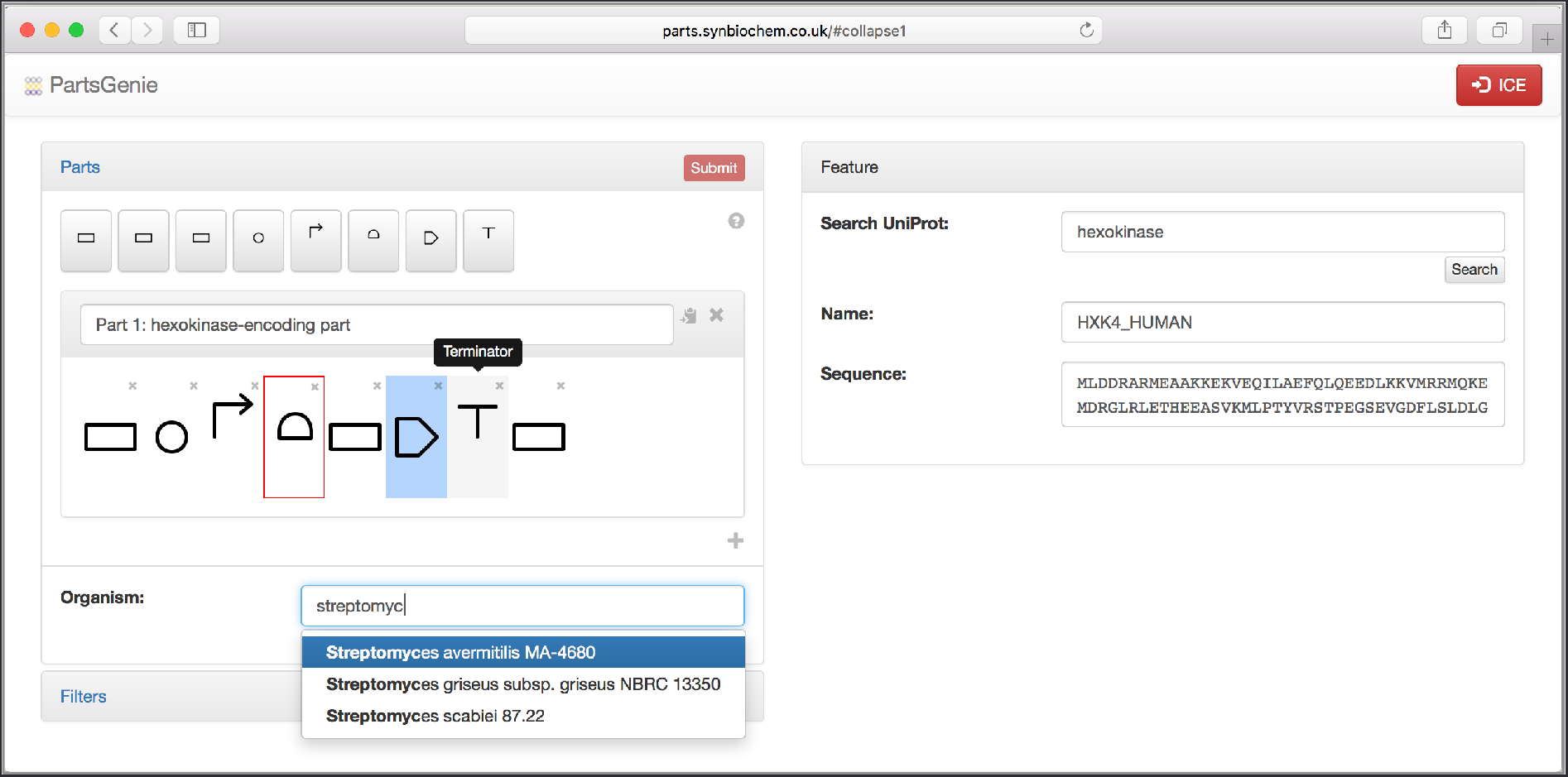
The PartsGenie drag-and-drop design interface. The Parts panel allows designs to be assembled from the palette of feature buttons above. Upon selecting a feature, the Feature panel allows for parameters necessary for the design and optimisation of that feature to be specified. Missing parameters and incorrectly ordered features (in this instance, an RBS not directly followed downstream by a CDS) are highlighted in red, and the job Submit button disabled until such errors are corrected, preventing the submission of invalid designs to the optimisation algorithm.

Filters can also be specified, which are applied to the design to prevent undesired subsequences appearing in the optimised sequence. These include limits on the number of repeating nucleotides, the exclusion of selected codons from CDS features, and the removal of unwanted restriction sites.

Upon submission, the user is presented with a progress panel, indicating the status of the running multi-objective optimisation, a simulating annealing algorithm that optimises all objectives simultaneously [10]. The progress panel indicates the codon adaptation index (CAI) [11] of any supplied CDS features, indicating codon suitability for the target host organism; a measure of the difference between desired and actual TIR for RBSs; the number of invalid sequences in the design, as defined by the filters; and the number of “rogue” RBSs, that is, sites that resemble RBSs with a sufficiently high TIR which may therefore reduce the efficiency of translation initiation at the intended RBS. Invalid sequences and rogue RBSs are removed from the design through optimisation before designed parts are returned.

Optimised designs can be viewed in an interactive Results panel (Figure 2), showing the result of each submitted design and allowing each individual feature to be investigated. Designs may be saved to any JBEI ICE repository, including the ACS Synthetic Biology Registry. PartsGenie therefore supports data sharing with standards such as SBOL, Genbank and fasta via its integration with ICE, from which these formats may be exported.

**Figure 2:**
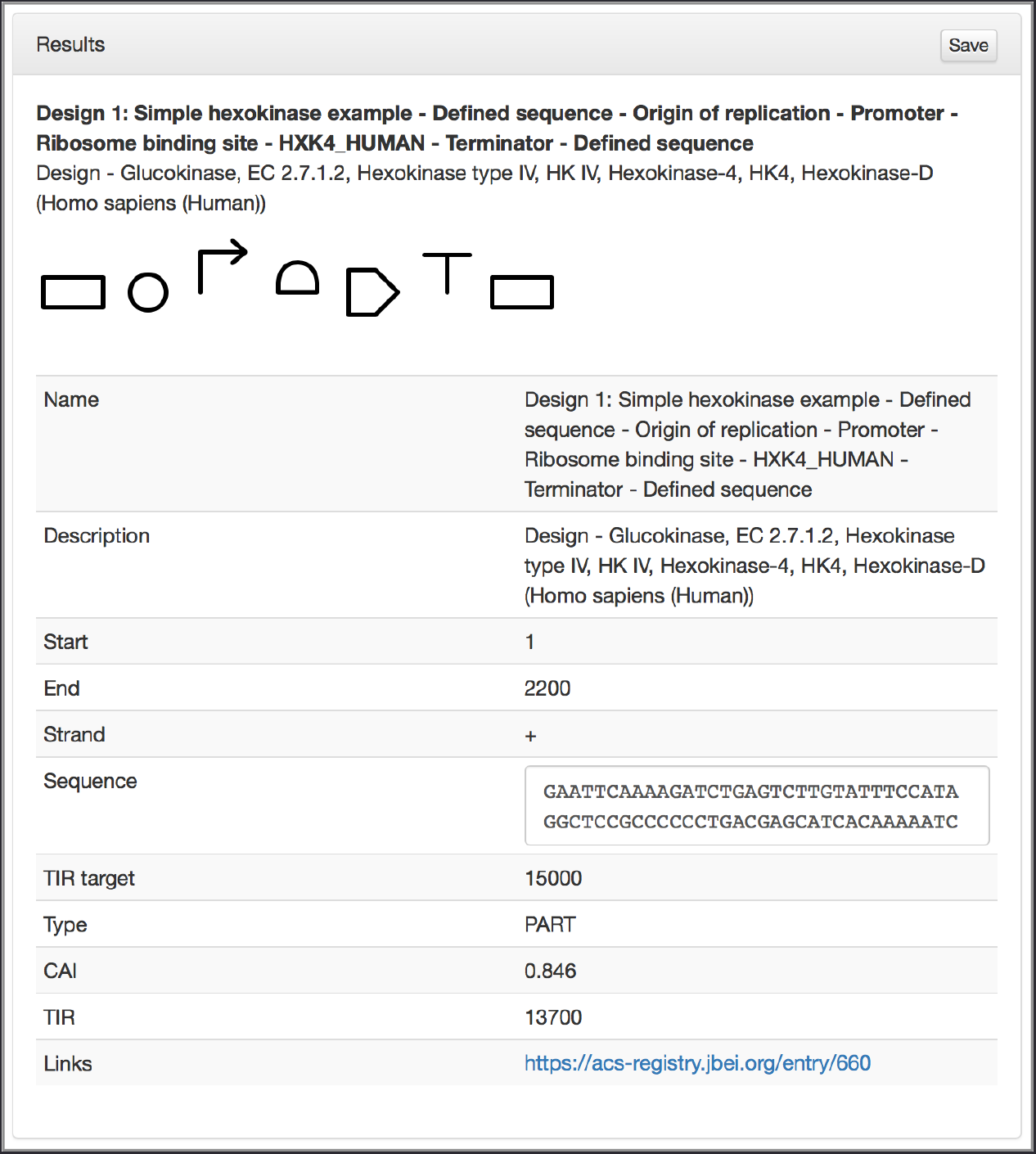
Results panel showing an optimised design. The whole design, including sequence and associated metadata is visible by default. Subsequence and metadata can be viewed by selecting a given feature within the design. Note that this example has been submitted to the ACS Synthetic Biology ICE repository, and a link to this entry is also provided.

PartsGenie has been designed to be sufficiently generic to be of use in a range of synthetic biology studies. However, we envisage that over time PartsGenie will be integrated into a larger computational design pipeline for metabolic engineering within the SYNBIOCHEM centre. Such a pipeline will include a number of existing and novel software tools, including the upstream modules RetroPath [12] for pathway elucidation and Selenzyme [13] for enzyme selection. Downstream of PartsGenie, tools for reducing experimental complexity through Design of Experiments approaches, and automated methods for the assembly of genetic constructs through interfacing with robotics liquid handling systems, will be introduced.

## Methods

PartsGenie is written as a two-tier, single-page web application, using Python, Javascript and the Bootstrap and AngularJS web development libraries. The multi-objective optimisation algorithm follows a simulated annealing approach. TIR values for RBSs are calculated using a performance-optimised version of the RBS Calculator, which relies on the NUPACK library [14] and caches calculated TIRs to prevent unnecessary and expensive recalculations. The system interfaces with ICE through its RESTful API (http://ice.jbei.org/api/). PartsGenie is hosted on Google Compute Engine and is distributed as a Docker file with source code openly available under MIT License at https://github.com/synbiochem/PathwayGenie.

## Supporting Information

An example PartsGenie part was submitted to the ACS Synthetic Biology Registry and is available at https://acs-registry.jbei.org/entry/660.

## Author Information

Authors’ address: BBSRC/EPSRC Manchester Centre for Synthetic Biology of Fine and Speciality Chemicals (SYNBIOCHEM), Manchester Institute of Biotechnology, The University of Manchester, Manchester M1 7DN, United Kingdom.

## Author Contribution

NS conceived and led the project, developed and tested the software, and led the writing of the paper. MD, AJ, CR and PC contributed suggested improvements and tested the software. PC and AW provided technical assistance. All authors contributed to writing of the paper.

## Acknowledgements

All authors acknowledge the funding from the Biotechnology and Biological Sciences Research Council (BBSRC) and the Engineering and Physical Sciences Research Council (EPSRC) under grant BB/M017702/1, “Centre for synthetic biology of fine and speciality chemicals (SYNBIOCHEM)”. NS is grateful for help received from James McLaughlin, Neil Wipat, Hector Plahar, Nathan Hillson and the SBOL development team.

This is a contribution from the Manchester Centre for Synthetic Biology of Fine and Speciality Chemicals (SYNBIOCHEM).

